# Tn*3*-derived inverted-repeat miniature elements (TIMEs) that mobilize antibiotic resistance genes

**DOI:** 10.1101/2025.11.05.686661

**Authors:** Ryota Gomi, Hirokazu Yano

## Abstract

Miniature inverted-repeat transposable elements (MITEs) are nonautonomous mobile genetic elements (MGEs) that can be mobilized by transposases provided by the relevant autonomous MGEs. MITEs originating from Tn*3*-family transposons were previously termed Tn*3*-derived inverted-repeat miniature elements (TIMEs). Composite transposon-like structures bounded by two copies of TIME, called TIME-COMPs, were shown to mobilize the intervening sequences. However, their association with antibiotic resistance genes (ARGs) has not yet been systematically studied. This study thus aimed to identify new TIME-COMP-like structures containing ARGs in the genomic sequences of the clinically important bacterial family Enterobacteriaceae in public databases. TIME-COMP-like structures were first searched for in the plasmid database PLSDB, focusing on small plasmids, using a self-against-self blastn approach to identify repeated elements. Then, newly and previously identified MITEs (including TIMEs) were searched for in the NCBI core nucleotide database to identify TIME-COMP-like structures located on other replicons. Bioinformatic analysis identified multiple previously unreported TIME-COMPs containing ARGs, which are bounded by directly or inversely oriented TIMEs, namely, IS*101*, MITESen1, and a novel 244-bp TIME termed TIME244. TIME244 contains a putative resolution site related to that of Tn*21*. These TIMEs were predominantly detected in plasmids and very rarely in chromosomes. The ARGs embedded in newly identified TIME-COMPs were *bla*_KPC-2_, *floR*, *qnrS1*, and *tet*(A). Notably, the *bla*_KPC-2_ carbapenemase gene was found in TIME-COMPs bounded by TIME244 and a TIME-COMP bounded by IS*101*. These findings highlight a potential role for TIMEs in the spread of diverse ARGs.

**IMPACT STATEMENT:** Bacterial miniature inverted-repeat transposable elements (MITEs) are a group of short (50 bp–500 bp) nonautonomous transposable elements that are thought to have originated from insertion sequences or transposons. Although MITEs can theoretically mobilize antibiotic resistance genes (ARGs) in the presence of transposases, only a few studies have reported their association with ARGs, probably due to difficulties in identifying MITEs in genomic sequences. This study provides evidence, based on bioinformatic analysis of public Enterobacteriaceae genomes, that a subset of MITEs, called Tn*3*-derived inverted-repeat miniature elements (TIMEs), mobilizes ARGs by forming composite transposon-like structures. A novel 244-bp TIME, designated TIME244, was present in more than 100 Enterobacteriaceae plasmids in the current RefSeq database, suggesting its further transmission in bacterial populations through horizontal gene transfer. This study reveals that TIMEs were often overlooked when analyzing the genetic contexts of ARGs in previous studies. These findings highlight the importance of TIMEs in bacterial gene acquisition and underscore the need for new tools that can detect TIMEs in bacterial genomes for ARG surveillance.

**DATA SUMMARY:** Accession numbers of sequence data analyzed in this study are provided within the article or in supplementary data files.

## INTRODUCTION

Antibiotic-resistant bacteria, particularly those with multidrug resistance (MDR), are a global health concern. Bacteria can accumulate antibiotic resistance genes (ARGs) with the help of mobile genetic elements (MGEs), which can often lead to MDR. MGEs can be categorized into two types: those mobilizing ARGs from one cell to another cell (i.e., intercellular mobilization) and those mobilizing ARGs from one replicon to another replicon or to different locations within the same replicon (i.e., intracellular mobilization).^1^ Intercellular mobilization mechanisms such as conjugation can directly contribute to horizontal transfer of ARGs and thus the emergence and spread of MDR. Intracellular mobilization can also contribute to the increase of MDR in multiple ways. For example, it can lead to the movement of ARGs from the chromosome to a plasmid, or from a plasmid to different plasmids, thereby increasing the potential of the ARGs to be transmitted to other bacterial recipients via conjugation.^2^ Additionally, it enables the accumulation of ARGs in a single genomic location to form a multiresistance region,^3^ facilitating their co-transfer as a single unit.

Transposable elements (TEs) can facilitate the intracellular mobilization of ARGs. TEs frequently associated with ARGs include insertion sequences (ISs), IS-composite transposons, and unit transposons such as the Tn*3* family.^1^ The majority of these elements encode DDE transposases.^4^ DDE transposases catalyze reactions such as the excision of a double-stranded TE bounded by inverted repeats (IRs) from its flanking DNA (for example, Tn*5*) or the nicking of the 3’ ends of a TE (for example, Tn*3*), followed by the transfer of the two 3’-OH ends to the two strands of a target site. Subsequent gap filling of the resulting strand-transfer product generates direct repeats (DRs) that flank the newly inserted TE, serving as a hallmark of the transposition event.^4^ A subset of TEs, called nonautonomous elements, can also mediate intracellular mobilization but require the transposase activity provided in *trans* by autonomous elements.^1^ Among the known nonautonomous elements are structures called miniature inverted-repeat transposable elements (MITEs).^5^ MITEs are thought to have originated from ISs or transposons. They retain the terminal IRs but lack the internal regions, including the transposase gene(s). They are typically 50 bp–500 bp and also create DRs upon insertion.^6^

A subset of MITEs originating from Tn*3*-family transposons and carrying Tn*3*-related IRs are called Tn*3*-derived inverted-repeat miniature elements (TIMEs), and they include elements such as IS*101*, integron mobilization unit (IMU), MITESen1, and TIME1–TIME4.^7–10^ Composite transposon-like structures bounded by two copies of TIME, known as TIME-COMPs, were shown to mobilize the intervening sequences when the transposase activity of autonomous Tn*3*-family elements was provided in *trans*.^7, 9^

However, to date, known TIME-COMPs carrying an ARG(s) are limited to IMU-bounded TIME-COMPs carrying a carbapenemase gene *bla*_GES-5_ and an IS*101*-bounded TIME-COMP carrying a quinolone resistance gene *qnrD1*, even in the clinically important bacterial family Enterobacteriaceae (**Figure S1**).^9, 11, 12^ MITEs other than TIMEs should also be capable of mobilizing an ARG when the ARG is embedded within a TIME-COMP-like structure; however, to our knowledge, no structures bounded by non-TIME MITEs have been reported in Enterobacteriaceae.

Tn*3*-family transposons have been identified in nearly all bacterial phyla and are classified into several subgroups (including Tn*3* and Tn*21*) based on the phylogeny of their transposases.^13^ These elements mobilize through a "paste-and-copy" mechanism involving two distinct steps: cointegrate formation and cointegrate resolution. The first step is mediated by the transposase (the *tnpA* gene product), IRs, and the host replication machinery, generating a fusion replicon in which the donor and target replicons are bridged by directly repeated copies of the transposon. In the second step, the cointegrate is resolved into two separate molecules, each carrying a transposon insertion. This resolution is typically mediated by the transposon-encoded resolvase (the *tnpR* gene product) and its cognate target site (*res*), or alternatively through homologous recombination between the duplicated transposon sequences. TIME and TIME-COMP elements are hypothesized to employ a similar "paste-and-copy" mechanism; however, whether these elements contain functional resolution sites remains unclear in most cases, with the exception of IS*101*, which possesses a functional *res* site.^14^

Transposition of an IMU-bounded TIME-COMP was previously demonstrated by providing a transposase of IS*Sod9* belonging to the Tn*21* subgroup.^9^ IS*101* cointegrate formation and resolution were demonstrated in the presence of Tn*1000* (also called gamma-delta) belonging to the Tn*3* subgroup.^14^ Known TIMEs and TIME-COMPs create 5-bp (or sometimes 6-bp) DRs upon transposition like Tn*3*-family transposons.^7, 9, 13^ Thus, past transposition of a TIME-COMP can be inferred from the presence of 5-bp or 6-bp DRs flanking the outermost IRs of the TIME-COMP.

MITEs (including TIMEs) tend to be overlooked in standard annotation pipelines due to their lack of transposase gene(s). Furthermore, as of January 2026, the number of MITE sequences registered in the ISFinder database remained at 65, with only three originating from Enterobacteriaceae.^15^ Thus, the contribution of MITEs to the intracellular mobility of ARGs is likely underestimated. Hence, this study aimed to identify previously overlooked TIME-COMP-like structures carrying ARGs in Enterobacteriaceae, by data-mining.

## METHODS

### Plasmid sequences

Plasmid sequences (n = 72,556) were downloaded from PLSDB (v. 2024_05_31_v2), which is a curated and non-redundant plasmid database sourced from the NCBI database.^16^ Plasmid sequences meeting the following criteria were extracted: (i) at least one ARG was detected by ABRicate (v1.0.1, https://github.com/tseemann/abricate) with the ncbi database;^17^ (ii) at least one plasmid replicon was detected by ABRicate (v1.0.1) with the “--db plasmidfinder” option^18^ (this was to exclude sequences that are not likely to be true plasmids and to be conservative; for example, NZ_CP107371 is composed almost entirely of ARGs and ISs/transposons, suggesting a non-plasmid circular element or a misassembled circular contig); (iii) the plasmid host is Enterobacteriaceae; and (iv) the length is 20,000 bp or shorter. Here, we focused on small plasmids because their small size allows manual curation of the results of bioinformatic analysis (described in detail below). Plasmid sequences meeting the above criteria (n = 1,007) were subjected to self-against-self blastn analysis as described below to detect repeated sequences that could be MITEs.

### Detection of TIME-COMP-like structures carrying ARGs

The BLAST+ (v2.16.0) program was used for similarity searches. The makeblastdb and blastn functions were used with default parameters. Plasmid sequences with self-against-self blastn hits meeting the following criteria were retained: (i) the alignment length is between 50 bp and 500 bp; (ii) the percentage of identical matches is 98 % or higher; (iii) a sequence longer than 500 bp is present between the aligned part of the query sequence and that of the subject sequence. These criteria were applied to detect repeat sequences (50 bp–500 bp) surrounding a stretch of sequence (>500 bp) in a plasmid (see **Figure S2** for the rationale for these criteria). To confirm that the 50-bp to 500-bp repeat sequences are actually MITEs, a self-against-self blastn search was performed for each repeat sequence to identify IRs, which are the signature of MITEs. The blastn search was performed by using a word size of six to increase the sensitivity, and the result was visualized using CLC Main Workbench 24 (Qiagen, Hilden, Germany) (**Figure S3**). The putative MITE sequences were checked for identity to any known elements in the ISfinder database.^15^ Only TIME-COMP-like structures flanked by DRs, which are typically created upon insertion of the entire structure, were considered for further analysis. This was to exclude cases where two MITEs were independently inserted and coincidentally formed a TIME-COMP-like structure. The lengths of DRs looked for were determined with reference to those of the related ISs or transposons. Plasmids with TIME-COMP-like structures carrying ARGs were annotated using ISfinder and the blastn web tool (https://blast.ncbi.nlm.nih.gov/Blast.cgi).

The MITE(s) found in the above approach, previously described TIMEs associated with ARGs in Enterobacteriaceae (i.e., IMU and IS*101*),^9, 11, 12^ and Enterobacteriaceae MITEs listed in the ISfinder database (i.e., MITEKpn1 and MITESen1, excluding MITEEc1, which is an enterobacterial repetitive intergenic consensus [ERIC] sequence and present in multiple copies on the chromosome of *Escherichia coli*),^15, 19, 20^ were searched for in the NCBI core nucleotide database (core_nt) using the blastn web tool (last accessed July 2025). Replicons with two or more blastn hits with ≥95 % identity and ≥95 % coverage were manually screened to detect additional TIME-COMP-like structures carrying an ARG in Enterobacteriaceae. This enabled identification of TIME-COMP-like structures in small plasmids not indexed in PLSDB, in longer plasmids, and also in chromosomes. (Note that we also performed the online blastn analysis described above for TIME1–TIME4, which were originally described in *Pseudomonas* spp.,^7^ and confirmed the absence of these elements in Enterobacteriaceae.)

Putative ancestral plasmids without insertion of TIME-COMP-like structures were identified by performing the online blastn analysis against the NCBI core_nt database, using a sequence, which was prepared by joining the sequences flanking the TIME-COMP-like structure minus one copy of the DR, as a query sequence (last accessed January 2026).

Sequence similarity among putative *res* regions of Tn*3*-family elements was evaluated using the SSEARCH program (ssearch36) from the FASTA package v.36.3.8i with default parameters,^21, 22^ querying a multi-FASTA file of putative *res* regions against itself. Smith–Waterman scores (S-W) above 100 were considered significant matches.

### Detection of MITEs in RefSeq Enterobacteriaceae genomes

To investigate the prevalence of MITEs that were found to be associated with ARGs, we downloaded RefSeq Enterobacteriaceae genomes with an assembly level of “complete genome” using the NCBI Datasets command-line tools (v18.9.0) in October 2025 (n = 11,572).^23^ MITE sequences were searched for in the retrieved genomes using makeblastdb and blastn in BLAST+ (v2.16.0) with default parameters. We used the same criteria as the online blastn search to define the presence of MITE sequences (i.e., ≥95 % identity and ≥95 % coverage).

## RESULTS AND DISCUSSION

### Identification of TIME-COMP-like structures carrying ARGs

First, we conducted self-against-self blastn analysis to identify TIME-COMP-like structures in a subset of small Enterobacteriaceae plasmids registered in the curated plasmid database, PLSDB. This relatively small dataset (n = 1,007) made manual curation of the blastn results feasible, which includes confirmation of the IRs in putative MITEs and DRs flanking the entire TIME-COMP-like structures. These procedures revealed TIME-COMP-like structures carrying ARGs in six plasmids (**Table S1**). All these structures are bounded by TIMEs and thus are TIME-COMPs. One of these plasmids is pKFu015_4, which was previously reported to carry the IS*101*-bounded TIME-COMP containing *qnrD1* (**Figure S1**).^12^ The remaining five plasmids contain the same TIME-COMP structure with *bla*_KPC-2_ and a truncated *bla*_TEM_ gene, which is bounded by two copies of a 244-bp TIME. This element was submitted to ISfinder and designated as MITEEc2;^15^ however, TIME244 was also registered as a synonym, which is used in this study to explicitly highlight that this element is indeed a TIME. pCHE-A and pCHE-A1, which were previously reported to contain IMU-bounded TIME-COMPs (**Figure S1**), were detected by the self-against-self blastn searches but were filtered out in manual curation, because the TIME-COMP structures in these plasmids are not flanked by DRs.^9,^

^11^ We also detected six other plasmids carrying TIME-COMP structures with ARGs but not flanked by DRs, all bounded by MITESen1 (see **Figure S4** for details). These TIME-COMP structures may have the potential to transpose as a unit; however, these cases were filtered out to restrict the dataset to plasmids showing direct evidence of transposition.

To identify additional TIME-COMP-like structures in replicons other than small plasmids indexed in PLSDB, we performed online blastn searches using TIME244 and previously reported MITEs, including those classified as TIMEs (IS*101*, IMU, and MITESen1) and one non-TIME MITE (MITEKpn1), as query sequences (see **Table S2** for the sequence of each MITE). This online blastn analysis identified 141 additional replicons carrying TIME-COMPs with ARGs. These included 136 plasmids and one chromosome carrying TIME244-bounded TIME-COMPs (most of which contain *tet*(A)), one plasmid carrying an IS*101*-bounded TIME-COMP, and three plasmids carrying MITESen1-bounded TIME-COMPs (**Table S1**).

The genetic contexts of TIME-COMPs identified using the above approaches were thoroughly inspected as described below.

### TIME244-bounded TIME-COMPs

As mentioned above, we detected TIME244-bounded TIME-COMPs in a total of 142 replicons (i.e., five identified by the self-against-self blastn analysis, and 137 identified by the online blastn analysis). Three main types of TIME244-bounded TIME-COMPs were identified.

One of them is a TIME244-bounded TIME-COMP carrying *tet*(A) detected in 133 replicons (**Table S1**). This TIME-COMP is bounded by inversely oriented TIME244 copies and contains part of Tn*1721*, an 11-kb Tn*3*-family element carrying tetracycline resistance determinant genes originally identified on an *E. coli* plasmid (**Figure 1a**).^24, 25^ This TIME-COMP was inserted in three unique locations with DRs of AGCAA (n = 114), TATAA (n = 18), and TACTT (n = 1) (**Figure 1a** and **Table S1**). This indicates that three independent transposition events of this TIME-COMP had occurred in the past. In 114 replicons with DRs of AGCAA, two types of the TIME-COMP were identified: one with an inverted internal structure (n = 52) and the other without (n = 62) (see below for detailed discussion). Furthermore, there were several indels within the TIME-COMP structure, two of which involved alterations of ARG contents (**Figure S5**, see **Table S1** for detailed description). Briefly, in a plasmid, a resistance region containing *bla*_TEM-1B_ and *bla*_CTX-M-206_ is inserted within the left TIME244 copy. In the other plasmid, the Tn*1721*-derived region is partially deleted and replaced by a truncated Tn*3*-like transposon and a truncated Tn*4401*-like transposon containing *bla*_KPC-2_ (i.e., a variant of non-Tn*4401* elements [NTE_KPC_], which contains only a portion of Tn*4401*, the most common *bla*_KPC_-containing mobile element belonging to the Tn*3* family^26^). In these cases, the DR sequences flanking the TIME-COMPs are the common ones (AGCAA); thus the indels were likely introduced after the insertion of the original TIME-COMP carrying *tet*(A). The outermost IRs within the altered TIME-COMP structures are intact, implying the feasibility of further transposition if the relevant transposase activity is provided in *trans* (**Figure S5**).

**Figure 1.**
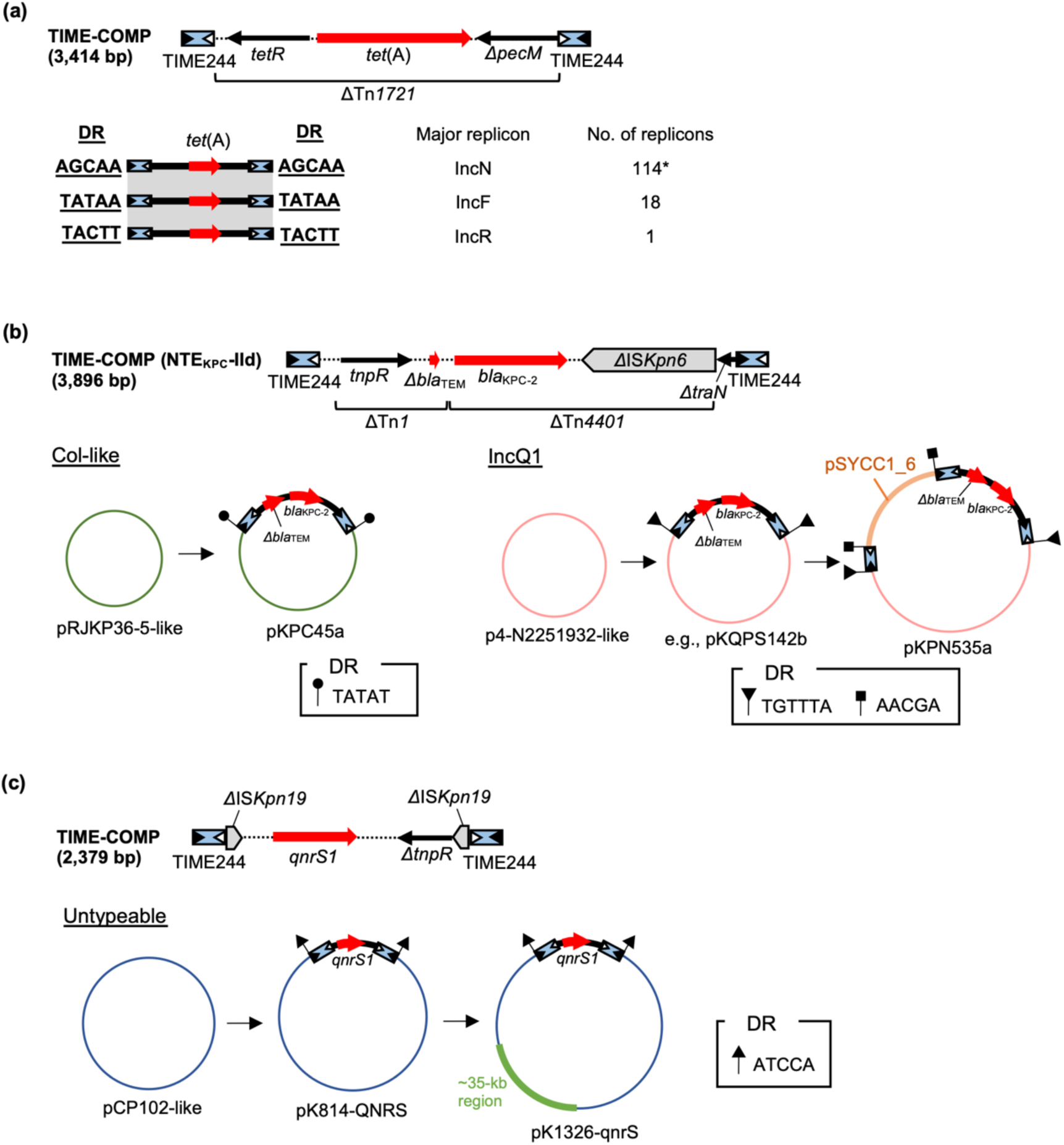
(a) TIME244-bounded TIME-COMP carrying *tet*(A) and DRs flanking the TIME-COMP. The expanded diagram of the TIME-COMP is shown on the top. Major replicon types and the numbers of replicons are shown on the right side. *There are variants in TIME-COMPs flanked by AGCAA (see **Table S1** for details). (b) TIME244-bounded TIME-COMP carrying *bla*_KPC-2_ and a truncated *bla*_TEM_ gene, and plasmids carrying the TIME-COMP. The expanded diagram of the TIME-COMP is shown above the plasmids. For the TIME-COMP structure in each plasmid, only TIMEs and ARGs are depicted for clarity. The TIME-COMP was inserted into a pRJKP36-5-like plasmid and a p4-N2251932-like plasmid, leading to pKPC45a and pKQPS142b (and four other IncQ plasmids showing >99% identity and >99% coverage against pKQPS142b, as in Table S1), respectively. In pKPN535a, three copies of TIME244 are present, and a sequence identical to pSYCC1_6 is present between two of the TIME244 copies. This indicates a cointegrate formation by “paste-and-copy” transposition of TIME244, leading to fusion of a pKQPS142b-like plasmid and pSYCC1_6. The online blastn analysis identified eight putative ancestral plasmids (including pRJKP36-5) for pKPC45 and seven putative ancestral plasmids (including p4-N2251932) for pKQPS142b, showing >90% identity and >98% coverage against the uninterrupted version of each plasmid. (c) TIME244-bounded TIME-COMP carrying *qnrS1*, and plasmids carrying the TIME-COMP. The TIME-COMP was likely inserted in a pCP102-like plasmid, leading to pK814-QNRS. pK814-QNRS then acquired a ∼35-kb region with high similarity to the partial sequence of *Klebsiella* phage KP12 clone KP12_2 (OM835952) (96% identity and 93% coverage), leading to pK1326-qnrS. The online blastn analysis identified three putative ancestral plasmids (including pCP102) for pK814-QNRS, showing >90% identity and >80% coverage against the uninterrupted version of pK814-QNRS. Red arrows indicate ARGs, gray pointed boxes indicate ISs, blue boxes indicate TIMEs, and black arrows indicate other genes. The IRs within each TIME are shown by a black (left, IRL) or white (right, IRR) triangle (see **Table S2** for the basis of the orientations of TIMEs). DRs are shown as differently shaped flags, and their sequences are shown in boxes. The accession numbers of the plasmids are as follows: pRJKP36-5, CP100081; pKPC45a, MH595534; p4-N2251932, CP165866; pKQPS142b, CP023480; pKPN535a, MH595533; pSYCC1_6, CP113183; pCP102, CP104015; pK814-QNRS, CP183858; and pK1326-qnrS, CP183864.

Another TIME-COMP carries *bla*_KPC-2_ and a truncated *bla*_TEM_ gene, which is bounded by directly oriented TIME244 copies (**Figure 1b**). This genetic context was previously described as a variant of NTE_KPC_, designated NTE_KPC_-IId.^27^ Although the study noted repeats at its boundaries, the size was described as 243 bp rather than 244 bp, likely because the Tn*3*-related IRs were not fully recognized. Furthermore, these repeats were not identified as MITEs or TIMEs. This TIME-COMP was inserted in the same site in six highly related IncQ1 plasmids, with the same DRs (TGTTTA), and in a single Col-like plasmid with DRs of TATAT (**Figure 1b** and **Table S1**). In one of the IncQ1 plasmids (pKPN535a), three copies of TIME244 are present, which could be explained by the fusion of two plasmids.

We also detected another TIME-COMP, which contains *qnrS1* and is bounded by inversely oriented TIME244 copies, in two untypeable plasmids (**Figure 1c**).^28, 29^ In these plasmids, the TIME-COMP structure is inserted in the same position with DRs of ATCCA; therefore, a total of one unique transposition event was detected. These two plasmids and their putative ancestral plasmid were classified as phages by geNomad (v1.11.2).^30^ Thus, these are putative phage-plasmids, which transfer horizontally between cells as viruses and vertically within cellular lineages as plasmids.^31^

### An IS*101*-bounded TIME-COMP

IS*101*, a 209-bp element, is among the earliest identified TIMEs and was originally discovered on a recombinant *E. coli* plasmid, pSC101.^8^ We identified a previously unreported IS*101*-bounded TIME-COMP in one *Enterobacter kobei* IncX3 plasmid, pEkFL23_IncX3 (**Figure 2**). This TIME-COMP is bounded by inversely oriented copies of IS*101* and carries *bla*_KPC-2_ embedded within a truncated Tn*4401b* structure.^32^ This genetic context was previously reported as Tn*4401k*, but the presence of IS*101* was not addressed in the previous study.^33^ The sequence between the two IS*101* copies in this TIME-COMP corresponds to the partial sequence of plasmids such as pKPC_FCF/3SP, and the whole TIME-COMP structure was flanked by 5-bp DRs (ATTCT). These indicate that IS*101*-mediated transposition of part of a pKPC_FCF/3SP-like plasmid into another plasmid generated pEkFL23_IncX3.

**Figure 2.**
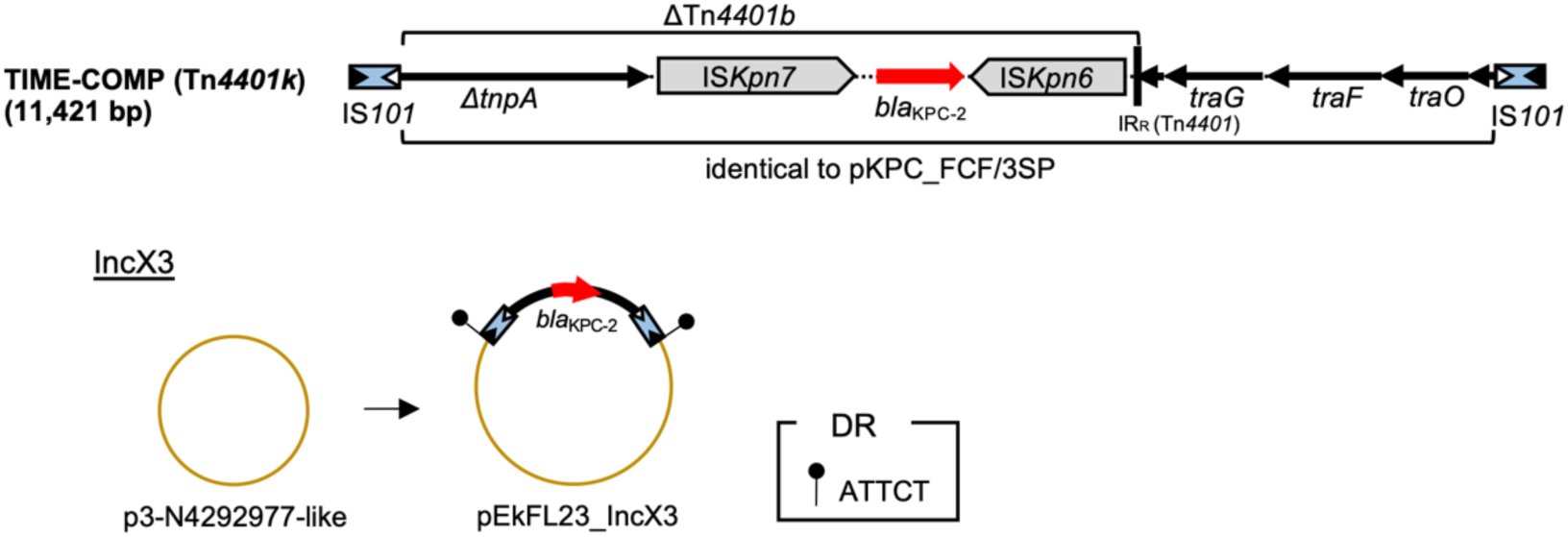
IS*101*-bounded TIME-COMP carrying *bla*_KPC-2_ and a plasmid carrying the TIME-COMP. The TIME-COMP was inserted into a p3-N4292977-like plasmid, leading to pEkFL23_IncX3. The online blastn analysis identified 19 putative ancestral plasmids (including p3-N4292977) for pEkFL23_IncX3, showing >99% identity and >90% coverage against the uninterrupted version of pEkFL23_IncX3. Genes and elements are shown as in Figure 1. The accession numbers of the plasmids are as follows: pKPC_FCF/3SP, CP004367; p3-N4292977, CP165819; and pEkFL23_IncX3, OQ434669.

### MITESen1-bounded TIME-COMPs

MITESen1 is a 256-bp TIME first identified on an IncX1 plasmid from *Salmonella enterica*.^10^ We identified two different MITESen1-bounded TIME-COMPs in the same location with the same flanking 5-bp DRs (TAATA) in two IncX1-IncX2 hybrid plasmids (**Figure 3a**).^34, 35^ One TIME-COMP contains *qnrS1*, and the other contains *floR*. MITESen1 copies are directly oriented in these TIME-COMPs. Interestingly, there are IncX1-X2 hybrid plasmids carrying a single MITESen1 copy inserted in the same location. We speculate that the two plasmids with TIME-COMPs were generated by integration of a hypothetical circular molecule containing MITESen1 and an ARG into a plasmid already carrying a single copy of MITESen1. The hypothetical circular molecules might be generated by recombination between MITESen1 in a TIME-COMP or by intramolecular replicative transposition of MITESen1 (also see **Figure S6** for another possible pathway for generation of the *floR*-containing circular element). Integration of the circular molecule might occur by homologous recombination between MITESen1 copies.

**Figure 3.**
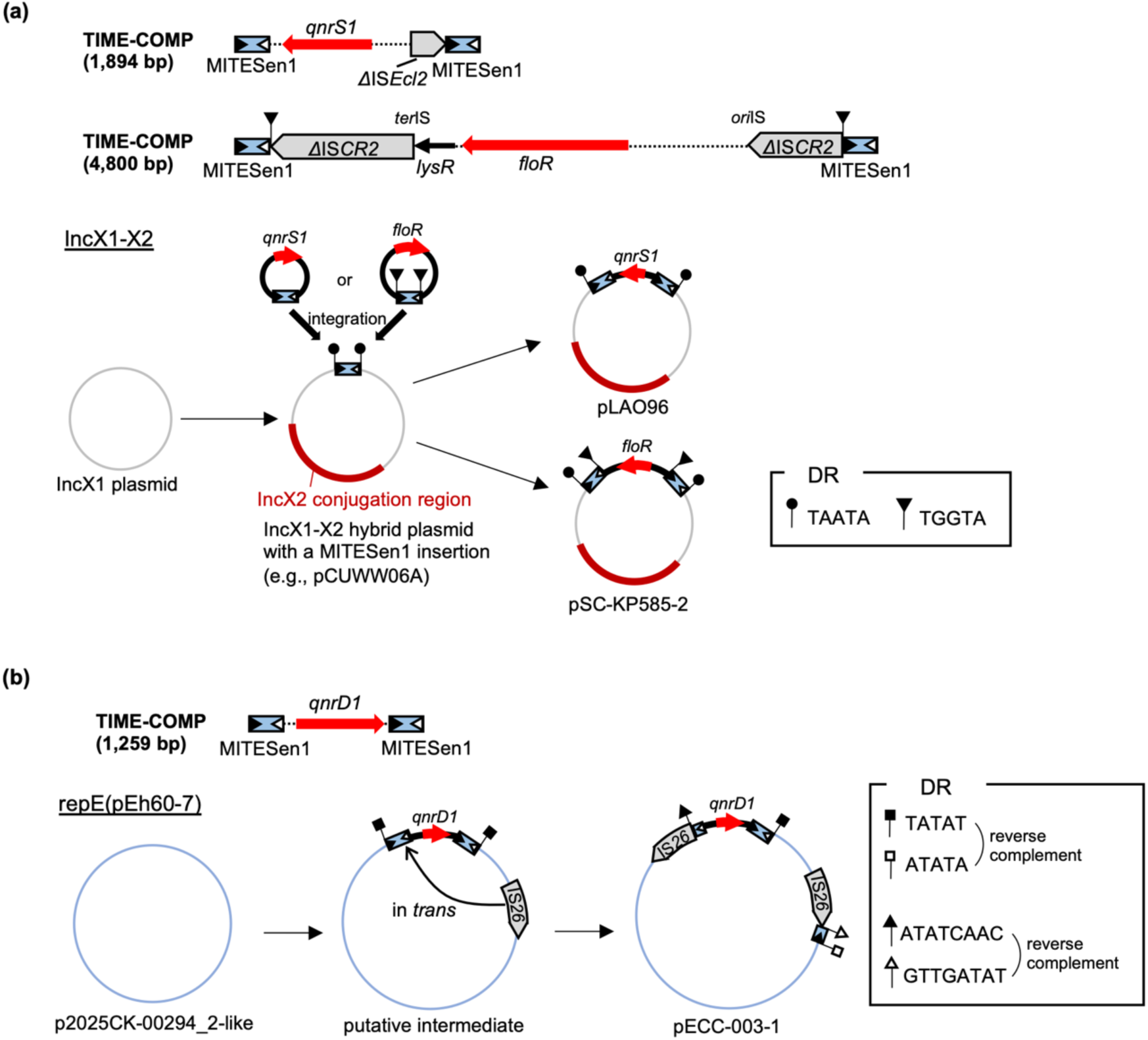
(a) MITESen1-bounded TIME-COMPs carrying *qnrS1* and *floR*, and plasmids carrying these TIME-COMPs. pLAO96 and pSC-KP585-2 might have been generated by integration of a hypothetical circular molecule into a pCUWW06A-like plasmid. pLAO96 (45,300 bp) and pSC-KP585-2 (44,092 bp) share ∼35.6 kb of highly similar sequence (>99% identity), excluding the TIME-COMP regions (1,894 bp and 4,800 bp) and regions bounded by IS*26* (7,824 bp and 3,609 bp). (b) MITESen1-bounded TIME-COMP carrying *qnrD1* and a plasmid carrying the interrupted TIME-COMP. A putative intermediate plasmid carrying the intact TIME-COMP is also shown. “in *trans*” indicates intramolecular replicative transposition (in *trans*) of IS*26*. The online blastn analysis identified one putative ancestral plasmid (p2025CK-00294_2) for pECC-003-1, showing >99% identity and 86% coverage against the uninterrupted version of the putative intermediate. Genes and elements are shown as in the previous figures. The accession numbers of the plasmids are as follows: pCUWW06A, CP182140; pLAO96, OP242301; pSC-KP585-2, CP123874; p2025CK-00294_2, CP194151; and pECC-003-1, CP143772.

A plasmid with the repE(pEh60-7) replicon (pECC-003-1) was found to carry a remnant of a MITESen1-bounded TIME-COMP carrying *qnrD1* (**Figure 3b**).^36^ One copy of MITESen1 is interrupted by IS*26*, while the truncated remnant of MITESen1 is present next to another copy of IS*26* that is oriented in the opposite direction. This arrangement is consistent with intramolecular replicative transposition (in *trans*) of an IS*26* element into MITESen1 in the intact TIME-COMP structure.^37^ The presence of 8-bp DRs (ATATCAAC or its reverse complement) next to each IS*26* element on the MITESen1-sides supports this, though the intact version of this TIME-COMP structure was not present in the NCBI core_nt database.

### Overview of TIME-COMP characteristics

**Table 1** summarizes the TIME-COMPs identified in this study as well as those reported previously. Four types of TIMEs (i.e., TIME244, IS*101*, MITESen1, and IMU) form TIME-COMPs with ARGs. Seven different intact ARGs were identified in these TIME-COMPs, including the carbapenemase genes *bla*_KPC-2_ and *bla*_GES-5_. Interestingly, the same TIMEs are associated with different ARGs, and the same ARGs are associated with different TIMEs. For example, TIME244 is associated with *tet*(A), *bla*_KPC-2_ (and Δ*bla*_TEM_), and *qnrS1*, while *bla*_KPC-2_ and *qnrS1* are also associated with IS*101* and MITESen1, respectively. The TIME-COMPs exhibit a broad size distribution, spanning from slightly above 1 kbp to over 10 kbp, implying that TIMEs can mediate the transposition of relatively long DNA segments. The replicon types carrying these TIME-COMPs are diverse; however, a distinct correspondence is observed between specific DR sequences and replicon types. This supports the idea that a TIME-COMP was inserted into an ancestor of that replicon type in a single event, followed by minor rearrangements that yielded the currently detected variants. However, in some cases for the TIME-COMP with *tet*(A), the TIME-COMP is carried by replicons different from the major replicon types (**Table S1**), indicating that MGEs other than TIMEs transposed a larger segment containing the TIME-COMP to other types of replicons. Of particular note is the overwhelming dominance of the TIME-COMP carrying *tet*(A), which may be attributed to two main reasons. One is the structural stability of the TIME-COMP bounded by inversely oriented TIMEs, as detailed below. The other is the successful spread of IncN plasmids carrying the TIME-COMP, as exemplified by the detection of these plasmids across multiple genera (i.e., *Citrobacter*, *Escherichia*, *Enterobacter*, *Klebsiella*, *Phytobacter*, and *Salmonella*; **Table S1**).

**Table 1.**
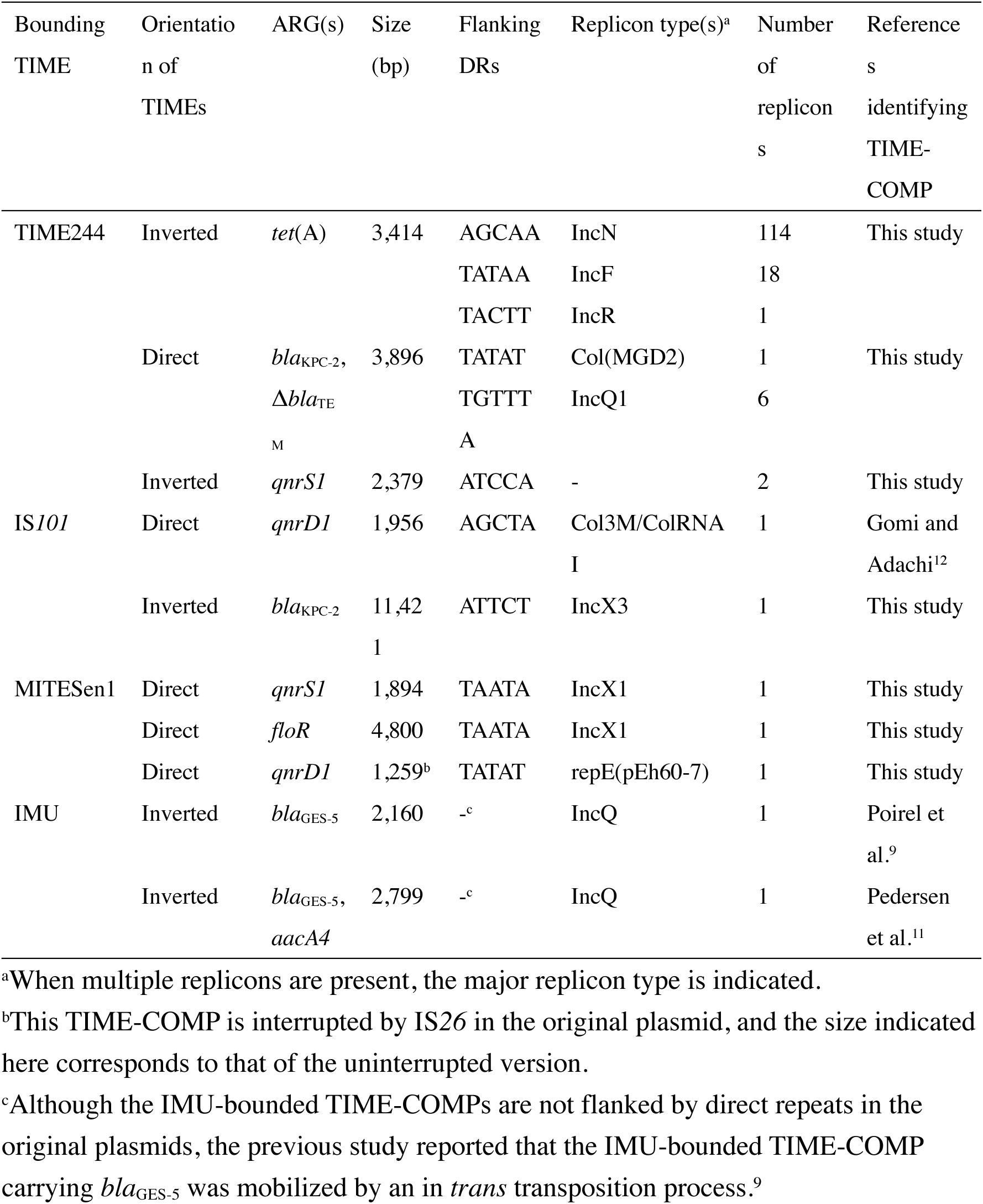
Summary of TIME-COMPs carrying ARGs in Enterobacteriaceae.

### Putative resolution sites in TIMEs

All TIMEs forming the TIME-COMP structures identified in the present study (i.e., TIME244, IS*101*, and MITESen1) carry IRs related to Tn*3*-family transposons (**Figure 4a**). IRs of TIME244 and the previously reported IMU show more similarities to IRs of Tn*21* than to those of Tn*3*, IS*101*, MITESen1, and *Pseudomonas*-derived TIME2 that is not associated with ARGs.^7^ Thus, TIME244 and IMU seem to have originated from the Tn*21* subgroup of the Tn*3* family.^13^ To deduce whether TIME244 contains a *res* site, we conducted all-against-all SSEARCH using sequences of TIMEs, Tn*21 res*, Tn*1721 res*, and Tn*3 res*.^22^ A significant match (Smith-Waterman score [S-W] >100) was detected for pairs of TIME244-Tn*21 res* (S-W = 148), TIME244-Tn*1721 res* (S-W = 152), Tn*21 res*-Tn*1721 res* (S-W = 188), and IS*101*-Tn*3 res* (S-W = 139). **Figure 4b** shows alignment of the TIME244 internal sequence and the experimentally determined *res* sites of Tn*21* and Tn*1721*.^38^ The *res* sites of the Tn*3* family consist of three subsites (site I, site II, and site III) where TnpR dimers bind. Strand exchange takes place at the center of site I (C/O in **Figure 4b**).^13^ The TIME244 internal sequence shows similarity to both Tn*21 res* and Tn*1721 res* at all three subsites. Therefore, we speculate that TIME244 contains a functional *res* site of the Tn*21* subgroup. On the other hand, the internal sequences of IMU and MITESen1 did not show similarity to well-characterized *res* sites from either autonomous Tn*3*-family elements or other TIMEs (S-W <70), although this does not exclude the possibility that these TIMEs carry resolution sites that serve as targets of site-specific recombinase(s).

**Figure 4.**
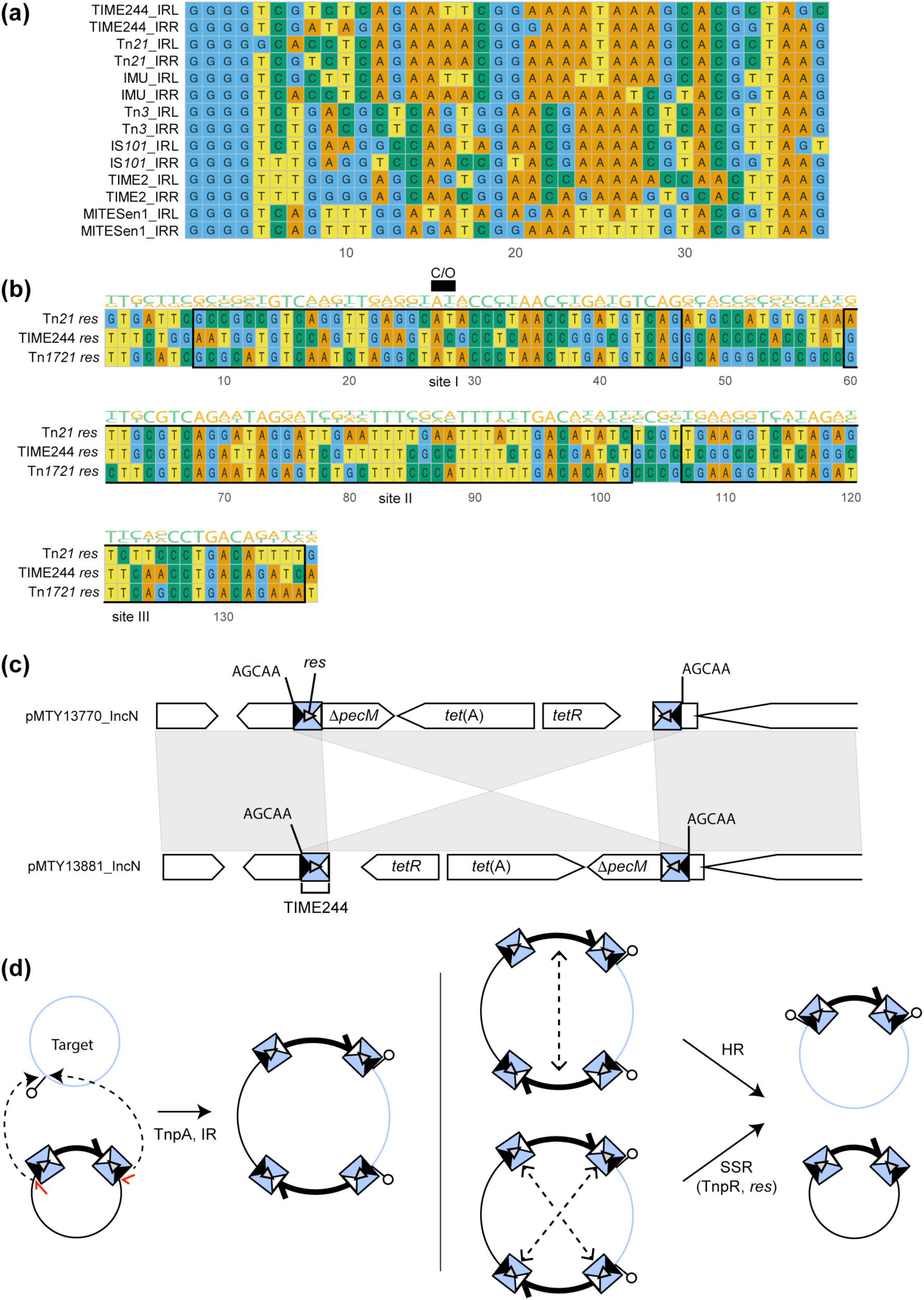
(a) Multiple alignment of IRs of TIMEs, Tn*3*, and Tn*21*. The alignment was visualized using the Bioconductor package ggmsa.^46^ (b) Multiple alignment of *res* sites of Tn*21*, TIME244, and Tn*1721*. Definition of subsites follows a previous study.^38^ See **Table S2** for the sequences of TIME244, IMU, IS*101*, and MITESen1. The remaining sequences are derived from the following accession numbers: Tn*21*, AF071413; Tn*3*, HM749966; TIME2, KJ920398; and Tn*1721*, X61367. Conservation level is indicated over the alignment by sequence logos in ggmsa. (c) Two frequent forms of TIME244-bounded TIME-COMPs carrying *tet*(A) inserted in the same location. The genetic contexts in two IncN plasmids (pMTY13770_IncN: LC720959 and pMTY13881_IncN: LC720960) are shown as examples.^47^ (d) Deduced transposition pathways of TIME-COMPs bounded by inversely oriented TIME244 copies. Cointegrate formation (left) is mediated by TnpA and IRs, followed by replication. Cointegrate resolution (right) is mediated by RecA-dependent homologous recombination (HR) between duplicated TIME-COMPs or by TnpR/*res*-dependent site-specific recombination (SSR). SSR can occur between either pair of the directly oriented TIME244 copies. Black circles indicate donor replicons, whereas light blue circles indicate target replicons. Filled and open triangles indicate the left and right terminal IRs of TIME244, respectively. Red arrows indicate nicking sites (3’ ends of TIME-COMP) in the donor molecule. Solid arrows indicate transition steps between structures. Dashed arrows connecting two regions indicate strand-nicking and transfer events between those regions, involved in cointegrate formation or resolution.

In principle, the functionality of *res* affects the stability of the TIME-COMP structures. When *res* sites are directly oriented, the segment between two *res* sites can be deleted in the presence of TnpR. According to studies on autonomous Tn*3*-family elements, when *res* sites are inversely oriented, the segment between the two *res* sites can be inverted by TnpR *in vivo*,^39, 40^ likely because knotted substrates carrying inversely oriented *res* sites, which are efficient substrates for TnpR,^41^ are produced *in vivo*. Among 142 replicons containing TIME244-bounded TIME-COMPs, only seven replicons contain directly oriented TIME244 copies, while 135 replicons contain inversely oriented TIME244 copies (**Table S1**). This orientation bias is consistent with the idea that TIME-COMPs carrying directly repeated TIME244 copies are structurally unstable due to the intrinsic resolution activity in the presence of Tn*21*-related TnpR.

The most prevalent TIME244-bounded TIME-COMP was that carrying *tet*(A). This TIME-COMP carries TIME244 copies in inverse orientation. The TIME-COMPs carrying *tet*(A) in the 114 replicons with DRs of AGCAA are classified into those with an inverted internal structure (n = 52) and those without (n = 62) (**Figure 1a** and **4c**). This inversion could be explained by TnpR-mediated site-specific recombination at the *res* sites. Because TIME244 contains a putative *res* site, the transposition of TIME244-bounded TIME-COMPs with inversely oriented TIME244 copies should follow a two-step process: cointegrate formation and cointegrate resolution in the presence of Tn*21*-related transposons (**Figure 4d**). Cointegrate resolution can occur by RecA-dependent homologous recombination or site-specific recombination involving Tn*21*-related TnpR and *res* sites.

### Prevalence of TIMEs in RefSeq Enterobacteriaceae genomes

The NCBI core_nt database, which we used in the online blastn analysis, contains a mixture of complete genomes and sequences that are not derived from complete genomes (e.g., plasmid sequences without the host chromosomal sequences). This prevents the accurate estimation of the prevalence of TIMEs associated with ARGs (i.e., TIME244, IS*101*, MITESen1, which were identified in this study within novel TIME-COMPs, as well as the previously reported IMU) in Enterobacteriaceae genomes. Thus, to circumvent this problem, we downloaded RefSeq complete Enterobacteriaceae genomes (n = 11,572) and detected these TIMEs in the retrieved genomes. The frequency of genomes carrying each TIME was as follows: TIME244 (n = 109, 0.94%), IS*101* (n = 49, 0.42%), MITESen1 (n = 264, 2.28%), and IMU (n = 7, 0.06%) (**Figure 5** and **Table S3**). The frequency of genomes with a replicon(s) carrying multiple copies of each TIME was as follows: TIME244 (n = 71, 0.61%), IS*101* (n = 4, 0.03%), MITESen1 (n = 54, 0.47%), and IMU (n = 2, 0.02%). These TIMEs were mainly detected in clinically important genera, namely *Klebsiella*, *Escherichia*, *Enterobacter*, and *Citrobacter*. However, these taxa are more likely to be sequenced due to their clinical importance, meaning the overall prevalence and burden of TIMEs in these genera may be inflated. Moreover, environmental or commensal bacteria, which are sequenced less frequently than clinical isolates, may harbor comparable TIME burdens. For example, the three *Leclercia* genomes carrying MITESen1 in our dataset were isolated from diverse sources (i.e., a clinical sample, a drain in a housekeeping closet in a hospital, and pig feed).^42–44^ Thus, care should be taken to interpret the results. In total, 408 genomes (3.53%) harbored at least one of these TIMEs, indicating that these elements, while rare, are present at a detectable frequency. In replicons carrying at least one TIME, 444 (97.16%) were plasmids and 13 (2.84%) were chromosomes.

**Figure 5.**
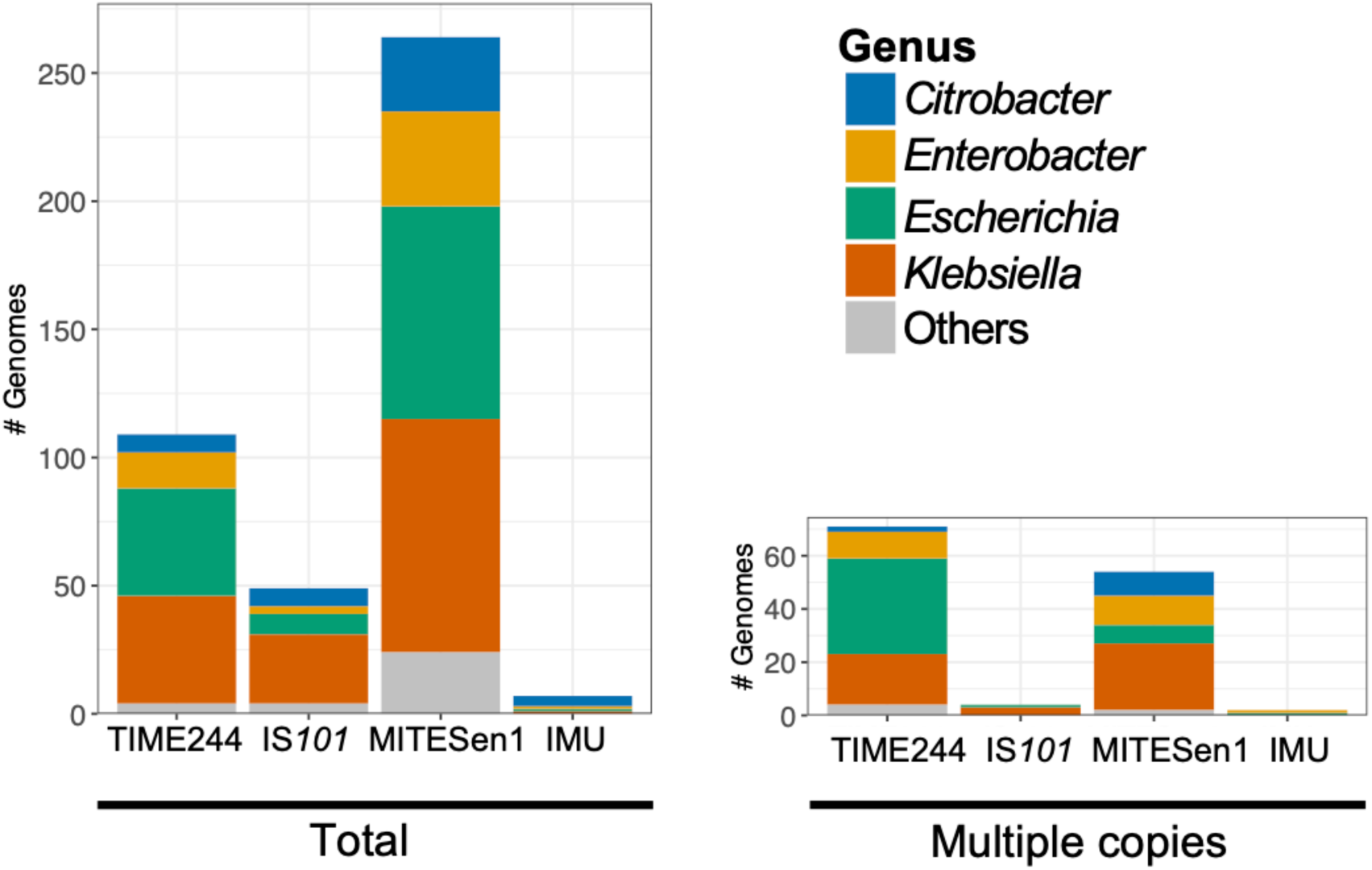
Numbers of RefSeq Enterobacteriaceae genomes carrying TIMEs (TIME244, IS*101*, MITESen1, and IMU) among the downloaded genomes (n = 11,572). “Total” indicates the total number of genomes carrying each TIME. “Multiple copies” indicates the number of genomes with a replicon(s) carrying multiple copies of each TIME. Genera detected in >10 genomes are shown in different colors, while genera detected in ≤10 genomes are grouped as “Others” and shown in gray. See also **Table S3** for information on individual genomes and **Table S4** for the numerical values presented in this figure.

### Study limitations

This study has limitations. First, the self-against-self blastn approach, which allows *de novo* identification of MITEs that form TIME-COMP-like structures, was performed only on small plasmids (≤20 kbp). Thus, novel MITEs potentially present in other types of replicons, such as larger plasmids and chromosomes, might have been missed. Specifically, there were 16,148 Enterobacteriaceae plasmids meeting criteria (i), (ii), and (iii) in the “Plasmid sequences” section; however, only 1,007 (6.24%) small plasmids (≤20 kbp) were targeted in our *de novo* identification analysis, leaving larger plasmids, including 1,872 (11.59%) in the 20–50 kbp range, unexamined. Performing the self-against-self blastn analysis also on larger replicons may enable the identification of more MITE elements; however, this approach is unwieldy because it will return exponentially more blastn hits, which makes manual curation of the results (e.g., confirmation of DRs at the ends) almost impossible. A direction for future work would be to develop software for automated detection of TIME-COMP-like structures. Moreover, there may be a prior literature bias in MITE discovery, which may have historically favored small plasmids due to the ease of sequencing. This bias may also impact the broader database searches, potentially underrepresenting MITEs unique to larger plasmids or chromosomes. A second limitation of this study is that we analyzed only the genomes of Enterobacteriaceae. Although rare, there seem to be TIME-COMP-like structures containing ARGs in other bacteria as well, such as *Acinetobacter* spp.^45^ Extending the approach employed in the present study to other bacteria may uncover additional TIME-COMP-like elements. Finally, while the transposition of certain TIME-COMPs was previously experimentally verified,^7, 9^ experimental validation of the TIME-COMPs identified in the present study (especially those bounded by TIME244, which could be tested by providing a Tn*21*-related transposase in *trans*) would be a highly valuable next step to complement our bioinformatic findings.

## Conclusions

The present study identified multiple previously unreported TIME-COMPs containing ARGs. These novel TIME-COMPs are bounded by three types of TIMEs (i.e., TIME244, IS*101*, and MITESen1). The sequences of these three TIMEs and also IMU are divergent and seem to have independently emerged from distinct transposons of the Tn*3* family. This study revealed that TIMEs contribute to the intracellular mobilization of ARGs and highlights the importance of taking these elements into account when analyzing the genetic contexts of ARGs. Accurate identification of not only autonomous MGEs but also nonautonomous MGEs, including TIMEs, will enable us to capture the mobility landscape of ARGs more comprehensively in genomic surveillance.

## Funding information

This work was supported by JSPS KAKENHI (grant number JP22K18038 to RG, JP25K01942 to HY) and the Environment Research and Technology Development Fund (JPMEERF20235R01 to RG) of the Environmental Restoration and Conservation Agency, provided by the Ministry of the Environment of Japan.

## Acknowledgments

We thank Dr. Naofumi Handa at University of California, Davis for helpful discussions. Computations were partially performed on the NIG supercomputer at ROIS National Institute of Genetics.

## Author contributions

Conceptualization: R.G. Methodology: R.G. Formal analysis: R.G. and H.Y. Data Curation: R.G. Visualization: R.G. and H.Y. Project administration: R.G. Funding acquisition: R.G. and H.Y. Writing - Original Draft: R.G. and H.Y. Writing - Review & Editing: R.G. and H.Y.

## Conflicts of interest

The authors declare that there are no conflicts of interest.

## Notes

### Competing Interest Statement

The authors have declared no competing interest.

### Summary of Updates

The main text, a table, figures, and supplementary material were updated.

## REFERENCES

1. Partridge SR, Kwong SM, Firth N, Jensen SO. Mobile genetic elements associated with antimicrobial resistance. Clin Microbiol Rev 2018; 31: e00088–17.

2. Lerminiaux NA, Cameron ADS. Horizontal transfer of antibiotic resistance genes in clinical environments. Can J Microbiol 2019; 65: 34–44.

3. Partridge SR. Analysis of antibiotic resistance regions in Gram-negative bacteria. FEMS Microbiol Rev 2011; 35: 820–55.

4. Curcio MJ, Derbyshire KM. The outs and ins of transposition: from mu to kangaroo. Nat Rev Mol Cell Biol 2003; 4: 865–77.

5. Delihas N. Impact of small repeat sequences on bacterial genome evolution. Genome Biol Evol 2011; 3: 959–73.

6. Minnick MF. Functional roles and genomic impact of miniature inverted-repeat transposable elements (MITEs) in prokaryotes. Genes (Basel) 2024; 15: 328.

7. Szuplewska M, Ludwiczak M, Lyzwa K et al. Mobility and generation of mosaic non-autonomous transposons by Tn*3*-derived inverted-repeat miniature elements (TIMEs). PLoS One 2014; 9: e105010.

8. Fischhoff DA, Vovis GF, Zinder ND. Organization of chimeras between filamentous bacteriophage f1 and plasmid pSC101. Journal of Molecular Biology 1980; 144: 247–65.

9. Poirel L, Carrer A, Pitout JD, Nordmann P. Integron mobilization unit as a source of mobility of antibiotic resistance genes. Antimicrob Agents Chemother 2009; 53: 2492–8.

10. Pong CH, Hall RM. An X1alpha plasmid from a *Salmonella enterica* serovar Ohio isolate carrying a novel IS*26*-bounded *tet*(C) pseudo-compound transposon. Plasmid 2021; 114: 102561.

11. Pedersen T, Sekyere JO, Govinden U et al. Spread of plasmid-encoded NDM-1 and GES-5 carbapenemases among extensively drug-resistant and pandrug-resistant clinical Enterobacteriaceae in Durban, South Africa. Antimicrob Agents Chemother 2018; 62: e02178–17.

12. Gomi R, Adachi F. Quinolone resistance genes *qnr*, *aac(6’)-Ib-cr*, *oqxAB*, and *qepA* in environmental *Escherichia coli*: insights into their genetic contexts from comparative genomics. Microb Ecol 2025; 88: 6.

13. Nicolas E, Lambin M, Dandoy D et al. The Tn*3*-family of replicative transposons. Microbiol Spectr 2015; 3: MDNA3-0060-2014.

14. Ishizaki K, Ohtsubo E. Cointegration and resolution mediated by IS*101* present in plasmid pSC101. Mol Gen Genet 1985; 199: 388–95.

15. Siguier P, Perochon J, Lestrade L et al. ISfinder: the reference centre for bacterial insertion sequences. Nucleic Acids Res 2006; 34: D32–6.

16. Molano LG, Hirsch P, Hannig M, et al. The PLSDB 2025 update: enhanced annotations and improved functionality for comprehensive plasmid research. Nucleic Acids Res 2025; 53: D189–D96.

17. Feldgarden M, Brover V, Gonzalez-Escalona N et al. AMRFinderPlus and the Reference Gene Catalog facilitate examination of the genomic links among antimicrobial resistance, stress response, and virulence. Sci Rep 2021; 11: 12728.

18. Carattoli A, Zankari E, Garcia-Fernandez A et al. *In silico* detection and typing of plasmids using PlasmidFinder and plasmid multilocus sequence typing. Antimicrob Agents Chemother 2014; 58: 3895–903.

19. Hulton CS, Higgins CF, Sharp PM. ERIC sequences: a novel family of repetitive elements in the genomes of *Escherichia coli*, *Salmonella typhimurium* and other enterobacteria. Mol Microbiol 1991; 5: 825–34.

20. Wilson LA, Sharp PM. Enterobacterial repetitive intergenic consensus (ERIC) sequences in *Escherichia coli*: Evolution and implications for ERIC-PCR. Mol Biol Evol 2006; 23: 1156–68.

21. Smith TF, Waterman MS. Identification of common molecular subsequences. J Mol Biol 1981; 147: 195–7.

22. Pearson WR. Searching protein sequence libraries: comparison of the sensitivity and selectivity of the Smith-Waterman and FASTA algorithms. Genomics 1991; 11: 635–50.

23. O’Leary NA, Cox E, Holmes JB et al. Exploring and retrieving sequence and metadata for species across the tree of life with NCBI Datasets. Sci Data 2024; 11: 732.

24. Mattes R, Burkardt HJ, Schmitt R. Repetition of tetracycline resistance determinant genes on R plasmid pRSD1 in *Escherichia coli*. Mol Gen Genet 1979; 168: 173–84.

25. Allmeier H, Cresnar B, Greck M, Schmitt R. Complete nucleotide sequence of Tn*1721*: gene organization and a novel gene product with features of a chemotaxis protein. Gene 1992; 111: 11–20.

26. Chen L, Mathema B, Chavda KD et al. Carbapenemase-producing *Klebsiella pneumoniae*: molecular and genetic decoding. Trends Microbiol 2014; 22: 686–96.

27. Cerdeira LT, Lam MMC, Wyres KL et al. Small IncQ1 and Col-like plasmids harboring *bla*_KPC-2_ and non-Tn*4401* elements (NTE_KPC_-IId) in high-risk lineages of *Klebsiella pneumoniae* CG258. Antimicrob Agents Chemother 2019; 63: e02140–18.

28. Luo X. Enterobacter hormaechei strain K814 plasmid pK814-QNRS, complete sequence. https://www.ncbi.nlm.nih.gov/nuccore/CP183858.

29. Luo X. Enterobacter hormaechei strain K1326 plasmid pK1326-qnrS, complete sequence. https://www.ncbi.nlm.nih.gov/nuccore/CP183864.

30. Camargo AP, Roux S, Schulz F et al. Identification of mobile genetic elements with geNomad. Nat Biotechnol 2024; 42: 1303–12.

31. Pfeifer E, Rocha EPC. Phage-plasmids promote recombination and emergence of phages and plasmids. Nat Commun 2024; 15: 1545.

32. Naas T, Cuzon G, Villegas MV et al. Genetic structures at the origin of acquisition of the beta-lactamase *bla*_KPC_ gene. Antimicrob Agents Chemother 2008; 52: 1257–63.

33. Guardatti RGM, Kraychete GB, Picao RC. Analysis of distinct *bla*_KPC_-encoding plasmids in an *Enterobacter kobei* strain recovered from recreational coastal water. Sci Total Environ 2024; 915: 169945.

34. Snaith AE, Dunn SJ, Moran RA et al. The highly diverse plasmid population found in *Escherichia coli* colonizing travellers to Laos and its role in antimicrobial resistance gene carriage. Microb Genom 2023; 9: 001000.

35. Kao CY, Kuo PY, Lin CC et al. Molecular characterization of colistin-resistant *Klebsiella pneumoniae* isolates and their conjugative *mcr*-carrying plasmids. J Infect Public Health 2024; 17: 102588.

36. Chen F-J, Liao Y-C, Liou C-H. Enterobacter cloacae strain ECC-003 plasmid pECC-003-1, complete sequence. https://www.ncbi.nlm.nih.gov/nuccore/CP143772.

37. He S, Hickman AB, Varani AM et al. Insertion sequence IS*26* reorganizes plasmids in clinically isolated multidrug-resistant bacteria by replicative transposition. mBio 2015; 6: e00762.

38. Rogowsky P, Halford SE, Schmitt R. Definition of three resolvase binding sites at the *res* loci of Tn*21* and Tn*1721*. EMBO J 1985; 4: 2135–41.

39. Altenbuchner J, Schmitt R. Transposon Tn*1721*: site-specific recombination generates deletions and inversions. Mol Gen Genet 1983; 190: 300–8.

40. Droge P, Cozzarelli NR. Recombination of knotted substrates by Tn*3* resolvase. Proc Natl Acad Sci U S A 1989; 86: 6062–6.

41. Yano H, Garruto CE, Sota M et al. Complete sequence determination combined with analysis of transposition/site-specific recombination events to explain genetic organization of IncP-7 TOL plasmid pWW53 and related mobile genetic elements. J Mol Biol 2007; 369: 11–26.

42. Sun Q, Wang H, Shu L et al. *Leclercia adecarboxylata* From Human Gut Flora Carries *mcr-4.3* and *bla*_IMP-4_-Bearing Plasmids. Front Microbiol 2019; 10: 2805.

43. Weingarten RA, Johnson RC, Conlan S et al. Genomic Analysis of Hospital Plumbing Reveals Diverse Reservoir of Bacterial Plasmids Conferring Carbapenem Resistance. mBio 2018; 9: e02011–17.

44. Mei CY, Jiao X, Wu H et al. Detection of *cfr* in *Leclercia adecarboxylata* from pig feed, China. J Antimicrob Chemother 2022; 77: 1500–2.

45. Zong Z. The complex genetic context of *bla*_PER-1_ flanked by miniature inverted-repeat transposable elements in *Acinetobacter johnsonii*. PLoS One 2014; 9: e90046.

46. Zhou L, Feng T, Xu S et al. ggmsa: a visual exploration tool for multiple sequence alignment and associated data. Brief Bioinform 2022; 23.

47. Ikegaya K, Aoki K, Komori K et al. Analysis of the stepwise acquisition of *bla*_CTX-M-2_ and subsequent acquisition of either *bla*_IMP-1_ or *bla*_IMP-6_ in highly conserved IncN-pST5 plasmids. JAC Antimicrob Resist 2023; 5: dlad106.

