## Supplemental Figures (S1-S6) and Supplementary Text for "Tn*3*-derived inverted-repeat miniature elements (TIMEs) that mobilize antibiotic resistance genes"

### Supplementary Figures

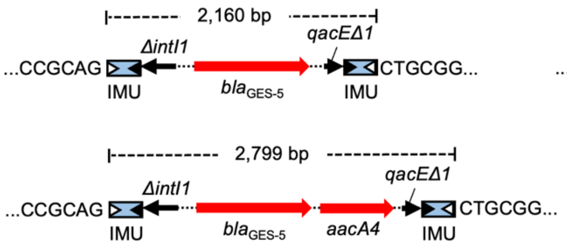


**Fig. S1.** Previously reported TIME-COMPs carrying ARGs found in Enterobacteriaceae. Red arrows indicate ARGs, blue boxes indicate TIMEs, and black arrows indicate other genes. The IRs within each TIME are shown by a black (left, IRL) or white (right, IRR) triangle. A TIME-COMP containing $bla_{GES-5}$ is from the *Enterobacter* sp. plasmid pCHE-A (NC_012006).[1] A TIME-COMP containing $bla_{GES-5}$ and *aacA4*, which is closely related to the TIME-COMP in pCHE-A, is from the *Klebsiella pneumoniae* plasmid pCHE-A1 (NZ_KX244760).[2] A TIME-COMP containing *qnrD1* is from the *Escherichia coli* plasmid pKFu015_4 (NZ_CP147134).[3] The 6-bp sequences flanking TIME-COMP structures are shown, and those corresponding to DRs are indicated in red. Only TIME-COMP(-like) structures showing direct evidence of transposition (i.e., those bounded by DRs or those with experimental evidence of transposition) are included and shown here. The IMU-bounded TIME-COMPs are not flanked by direct repeats in the original plasmids but are included here, because the previous study reported that the IMU-bounded TIME-COMP found in pCHE-A was mobilized by an in *trans* transposition process.[1]

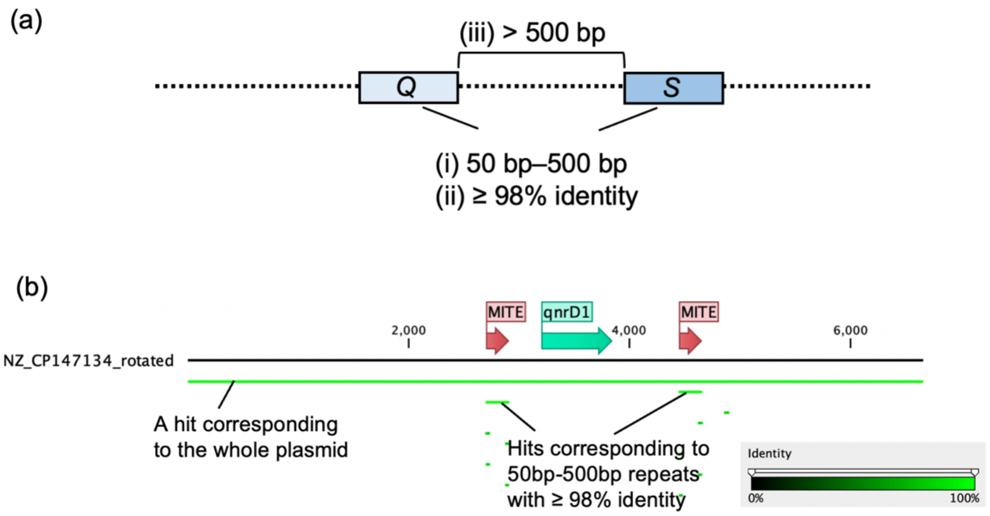


**Fig. S2**. (a) The strategy used to detect a potential TIME-COMP-like structure in a plasmid sequence by a self-against-self blastn search. *Q* indicates the aligned part of the query sequence, and *S* indicates the aligned part of the subject sequence. Note that *Q* and *S* are repeat sequences and not necessarily MITEs, and further analysis is needed to confirm that these are actually MITEs. Criteria (i)–(iii) were applied because (i) MITEs are typically 50 bp–500 bp in length,[4] (ii) previously reported TIME-COMP structures carrying ARGs are bounded by identical TIMEs,[1-3] and (iii) most ARGs reported to date are > 500 bp in length.[5] (b) An example of a self-against-self blastn search result for the *E. coli* plasmid pKFu015_4 (NZ_CP147134) carrying a TIME-COMP with *qnrD1*. The result was visualized with CLC Main Workbench 24. The plasmid sequence was rotated for visualization.

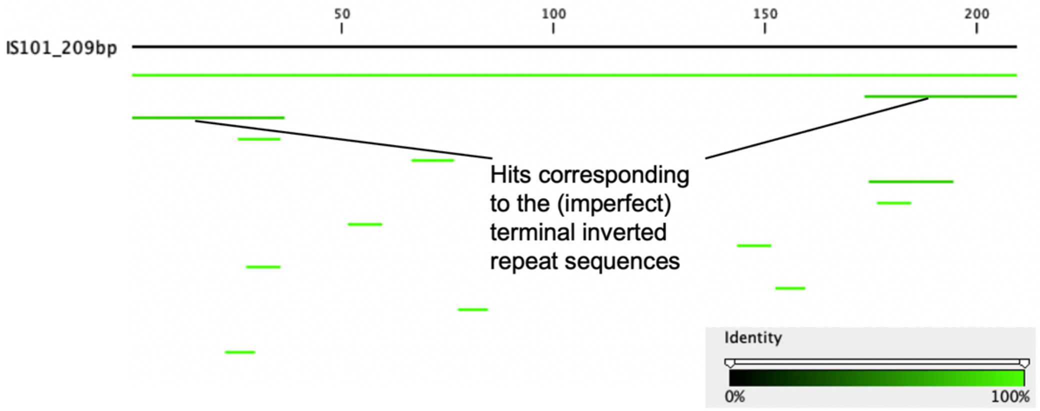


**Fig. S3**. An example of the result of a self-against-self blastn search of the repeat sequence. If the repeat sequence is actually a MITE, as in this figure (i.e., IS*101*), blastn hits with high nucleotide identity, which correspond to the (imperfect) terminal IRs, are present at each end of the sequence.

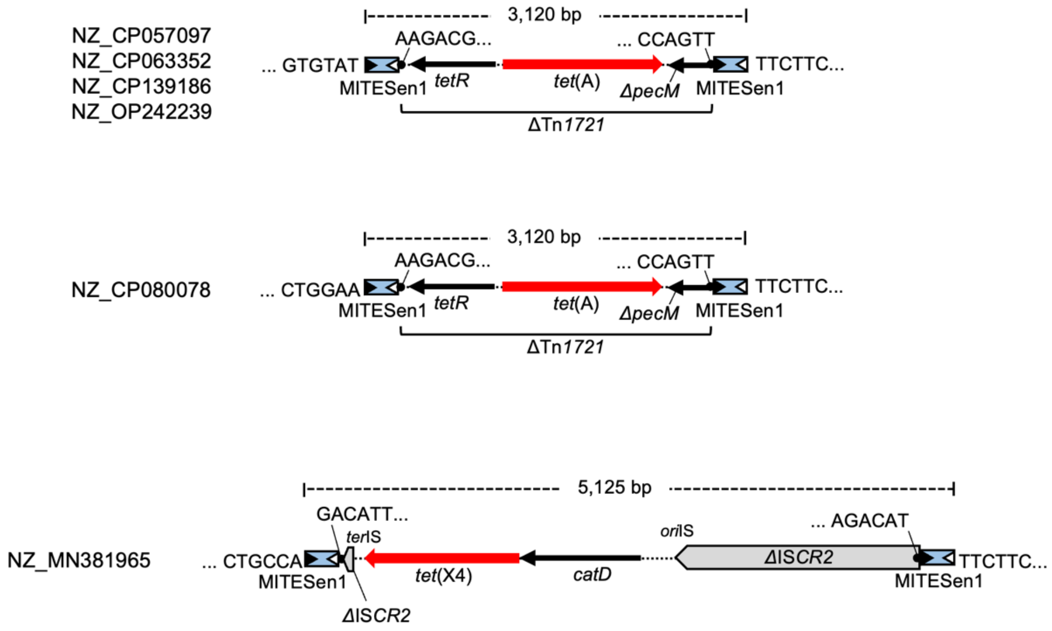


**Fig. S4**. Structures of TIME-COMPs without flanking DRs found by the self-against-self blastn analysis. GenBank accessions are shown on the figure. The 6-bp sequences flanking MITESen1 elements are shown. None of the TIME-COMP structures or the MITESen1 elements were bounded by DRs. These structures might have been formed by multiple independent insertions of the TIME-COMP structure or MITESen1 next to ARGs, followed by deletion of a segment between a copy of MITESen1 outside of the TIME-COMP structure and another copy in the TIME-COMP structure.

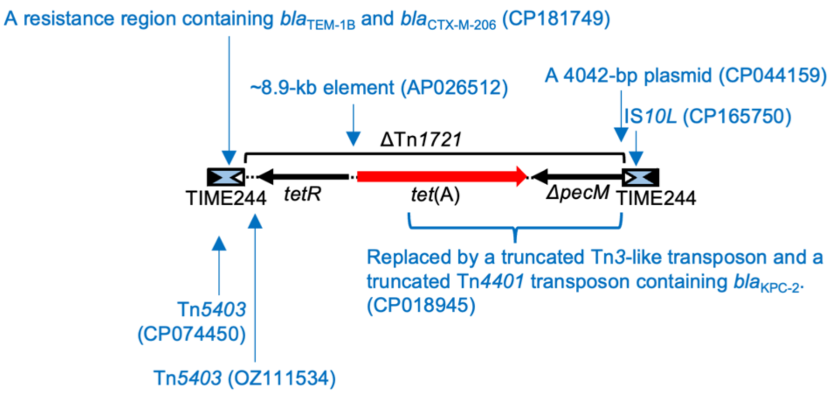


**Fig. S5.** Variants of the 3,414-bp TIME-COMP carrying *tet*(A). Insertion sites of MGEs are shown by vertical arrows, and the replaced region is shown with a curly bracket. GenBank accessions are shown on the figure.

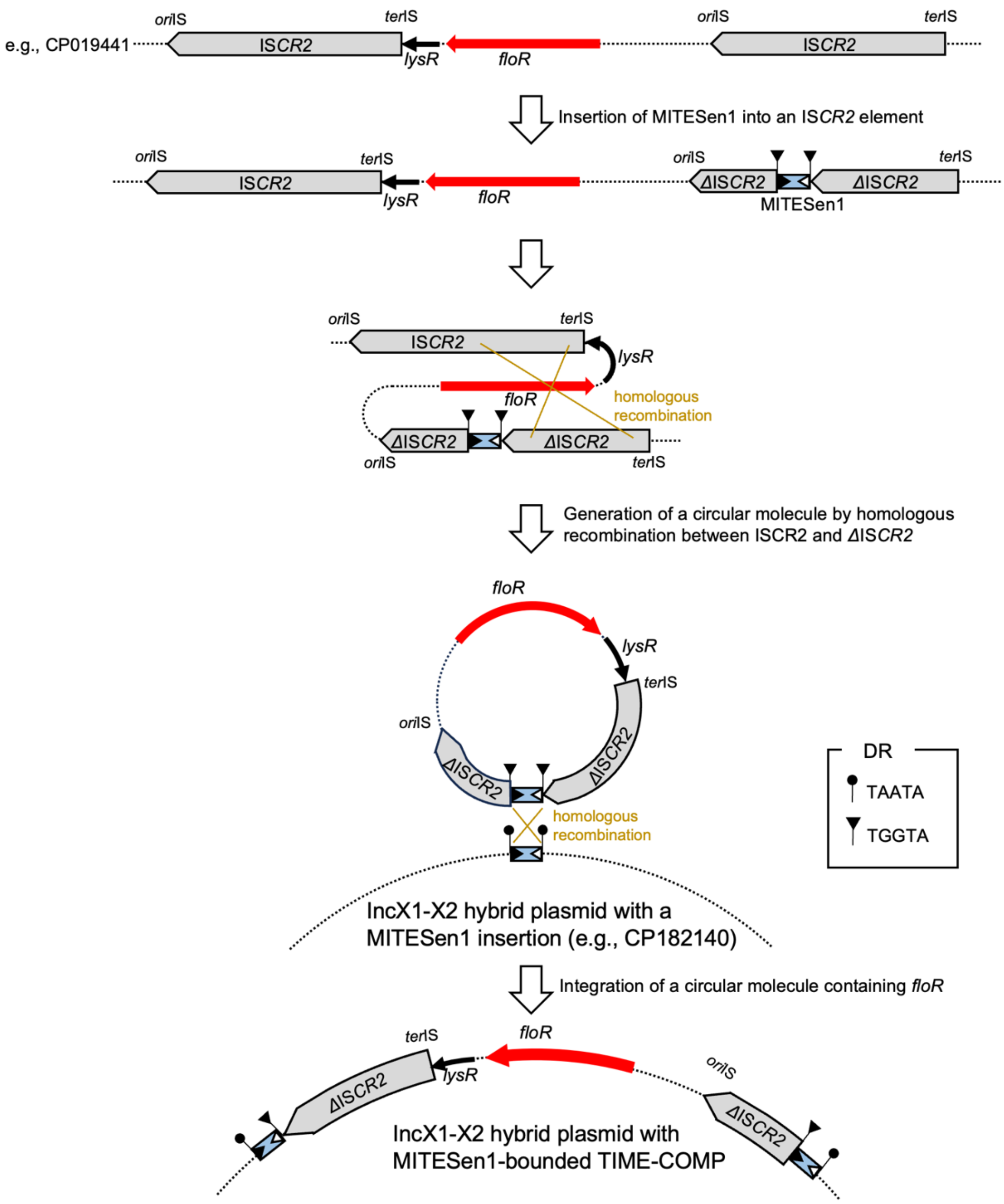


**Fig. S6**. A possible pathway for the formation of the genetic context of *floR* seen in pSC-KP585-2. A MITESen1 element was inserted into an IS*CR2*-*lysR*-*floR*-IS*CR2* element, which is present in replicons such as pETW41 (CP019441). This was followed by homologous recombination between IS*CR2* and ΔIS*CR2*, forming a circular molecule. This circular molecule was integrated into an IncX1-X2 hybrid plasmid possibly through homologous recombination between MITESen1 copies, generating the context seen in pSC-KP585-2.

**Supplementary Text**

Commands used for the ABRicate analysis

```
abricate --db ncbi --nopath --minid 80 --mincov 40 *.fasta >
abricate_args_out.tsv
#Go to the directory with FASTA files of interest and execute the command
above. This command detects ARGs using the ncbi database. We used a minimum
DNA %identity of 80% and a minimum DNA %coverage of 40% for sensitive
detection. Specifically, the minimum DNA %coverage of 40% allows detection of
ARGs that span the start and end of a contig.

abricate --db plasmidfinder --nopath --minid 80 --mincov 40 *.fasta >
abricate_args_replicons_out.tsv
#Run the above command on the FASTA files in which ARGs were detected. This
command detects plasmid replicons using the plasmidfinder database. We used
the same thresholds as those used for the detection of ARGs for the same
reason.
```

Commands used for the self-against-self blastn analysis to detect repeat sequences in small plasmids

```
for individual_file in *.fasta
do
 makeblastdb -in "$individual_file" -dbtype nucl -parse_seqids
 blastn -db "$individual_file" -query "$individual_file" -outfmt 6 >>
blastn_output.txt
done
#Go to the directory with FASTA files of interest and execute the command
above. Regarding the self-against-self blastn analysis for the identification
of IRs in each repeat sequence, the BLAST tool in CLC Main Workbench 24 was
used with a word size of six.
```

Commands used for running SSEARCH

```
ssearch36 TIME_res_sequences.fas TIME_res_sequences.fas > ssearch_results.txt
#TIME_res_sequences.fas is a multi-FASTA file of putative res regions.
```

Commands used for running geNomad

```
while read individual_file; do genomad end-to-end "$individual_file"
genomad_output_"${individual_file/.fasta}" /path/to/genomad_db; done <
fasta_list.txt
# Go to the directory with FASTA files of interest and execute the command
above. In fasta_list.txt, each line contains one FASTA file name.
```

Note that copying and pasting code directly from this file may introduce formatting or encoding errors; therefore, it is recommended to verify the characters or retype the command manually.

### REFERENCES

1. Poirel L, Carrer A, Pitout JD, Nordmann P. Integron mobilization unit as a source of mobility of antibiotic resistance genes. *Antimicrob Agents Chemother* 2009; **53**: 2492-8.

2. Pedersen T, Sekyere JO, Govinden U et al. Spread of plasmid-encoded NDM-1 and GES-5 carbapenemases among extensively drug-resistant and pandrug-resistant clinical Enterobacteriaceae in Durban, South Africa. *Antimicrob Agents Chemother* 2018; **62**: e02178-17.

3. Gomi R, Adachi F. Quinolone resistance genes *qnr*, *aac(6')-Ib-cr*, *oqxAB*, and *qepA* in environmental *Escherichia coli*: insights into their genetic contexts from comparative genomics. *Microb Ecol* 2025; **88**: 6.

4. Minnick MF. Functional roles and genomic impact of miniature inverted-repeat transposable elements (MITEs) in prokaryotes. *Genes (Basel)* 2024; **15**: 328.

5. Bortolaia V, Kaas RS, Ruppe E et al. ResFinder 4.0 for predictions of phenotypes from genotypes. *J Antimicrob Chemother* 2020; **75**: 3491-500.
